# Cannabis Use by People with HIV is Associated with an Anti-Inflammatory Immunometabolic Phenotype in Monocyte-Derived Macrophages

**DOI:** 10.64898/2026.03.04.709579

**Authors:** Mary K. Ford, Peter W. Halcrow, Anna E. Laird, Bryant L. Avalos, Ali Boustani, Matthew Spencer, Jack I. Melcher, Kyle C. Walter, Derek D. Hong, Gail Funk, Elizabeth G. Searson, Alexandra A. Le, Ronald J. Ellis, Scott Letendre, Maria Cecilia Garibaldi Marcondes, Johannes Schlachetzki, Jennifer Iudicello, Jerel A. Fields

## Abstract

Chronic neuroinflammation is associated with comorbidities in people with HIV (PWH) on antiretroviral therapy (ART). While cannabis use is associated with reduced neuroinflammation and neurocognitive impairment (NCI) in PWH, the underlying mechanisms are unknown. To address this gap in knowledge, we analyzed monocyte-derived macrophages (MDMs) from a cohort of 50 PWH and 33 people without HIV (mean age: 61.9 years), categorized by frequency of cannabis use (naïve/low, moderate, daily). We performed immunocytochemistry, RNA sequencing, and qPCR on MDMs and quantified related biomarkers in donor plasma. In this cohort study, daily cannabis use in PWH was associated with less global neurocognitive deficits, and with an anti-inflammatory immunometabolic-phenotype in MDMs characterized by (1) a metabolic shift from glycolysis to oxidative phosphorylation, (2) higher mitochondrial numbers, (3) altered cytokine profiles (pro-inflammatory downregulation, anti-inflammatory upregulation), and (4) higher brain-derived neurotrophic factor (BDNF) expression. These cellular changes were corroborated by a plasma biomarker profile in PWH including (1) lower levels of growth differentiation factor 15 and soluble triggering receptor expressed on myeloid cells 2, and (2) higher mature BDNF/precursor BDNF ratios that correlated with better cognition. Thus, cannabis use may mitigate NCI in PWH by immunometabolically reprogramming MDM function towards an anti-inflammatory and neuroprotective state.

**Graphical Abstract:** 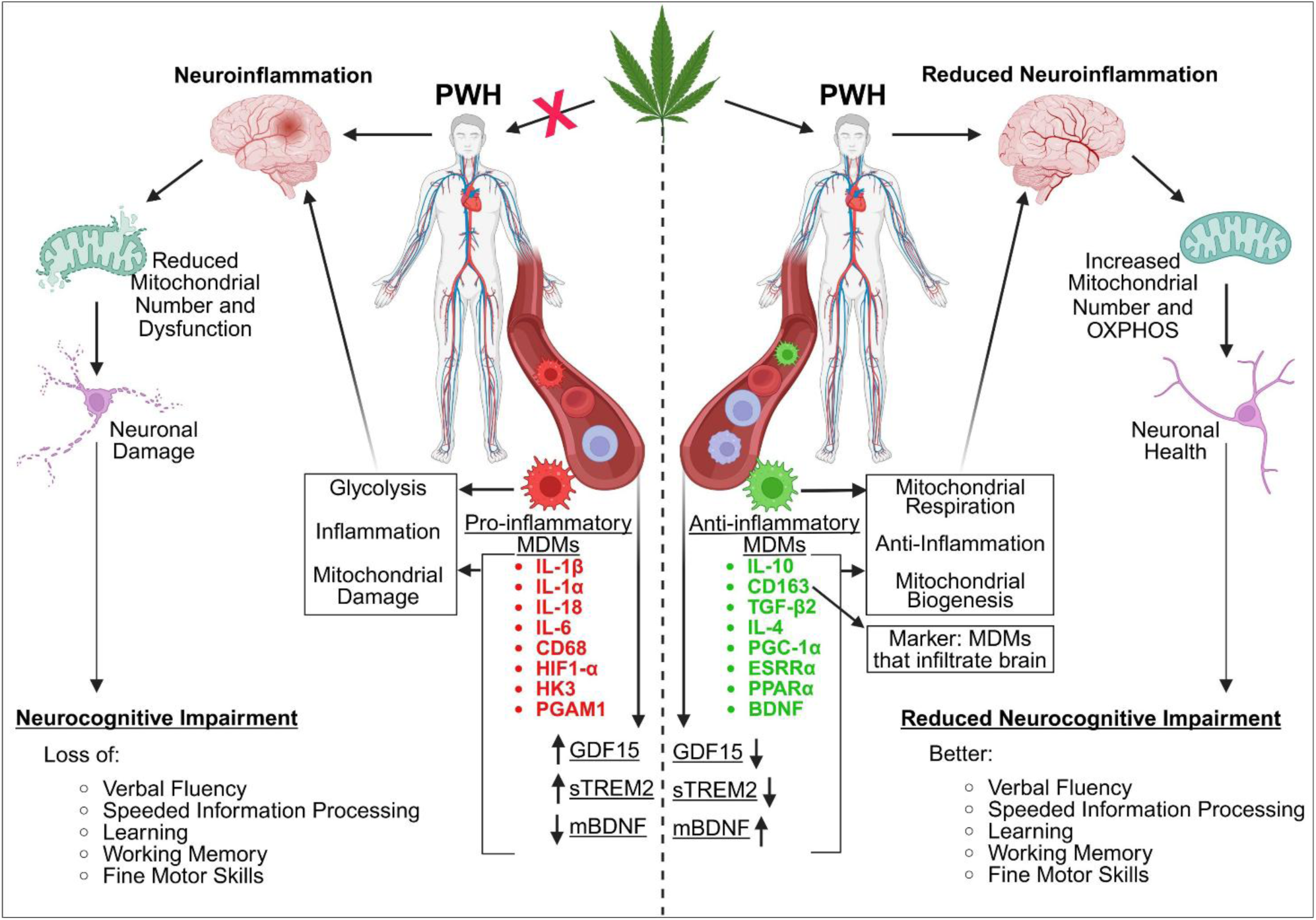

## Introduction

Antiretroviral therapy (ART) has transformed HIV into a manageable chronic condition, increasing the life expectancy of people with HIV (PWH). However, this advancement has revealed a new challenge: an aging population of PWH facing a range of comorbidities, driven by the virus’s long-term effects, particularly chronic neuroinflammation (1–4). This persistent inflammation can lead to HIV-associated neurocognitive impairment (NCI), which manifests as deficits in memory, learning, and motor function (5). These cognitive deficits contribute to morbidity and mortality among aging PWH on ART (6,7). Despite progress in understanding the pathogenic mechanisms of HIV-NCI, effective interventions are needed to preserve brain function in PWH.

Monocyte-derived macrophages (MDMs) are crucial for inflammatory responses, expressing both pro- and anti-inflammatory signaling proteins (8–10). Circulating monocytes migrate into tissues for purposes of routine surveillance in response to chemokines and extracellular stimuli, where they differentiate into pro- or anti-inflammatory macrophage phenotypes (8,9,11–13). Upon migrating into tissues, MDM polarization influences the local inflammatory milieu, particularly in the brain where it may affect neuronal function (14). While pro-inflammatory MDMs produce neurotoxic cytokines (e.g., TNF-α, IL-1β, IL-6) that can damage neurons, anti-inflammatory MDMs express factors like IL-10 and IL-4 that may promote neuroprotection and homeostasis (14–19). Although PWH on suppressive ART exhibit chronically elevated inflammatory markers in the blood and low-level neuroinflammation, the relationship between MDM phenotype and these inflammatory mediators is not well understood (20).

With its long history as a medicinal plant, cannabis contains components with potential therapeutic benefits (21). PWH use cannabis for both recreational and symptomatic relief at a rate two to three times higher than the general population (22–25). This use is associated with reduced inflammatory biomarkers in PWH and, in some instances, a lower prevalence of NCI (26–33). For example, studies have observed lower plasma levels of inflammatory markers like IL-16 and IP-10 in cannabis-using PWH (26,34). Despite evidence that cannabinoids can influence the inflammatory phenotype of MDMs, the specific cannabis use patterns and underlying mechanisms linking cannabis to less NCI in the context of HIV infection are not yet fully understood.

Cannabinoids may modulate several potential etiologies contributing to HIV-NCI, such as immune dysregulation, premature aging, and metabolic dysfunction (35–39). Even with suppressed viral loads, ART-treated PWH are susceptible to NCI. Postmortem studies of such individuals have revealed synaptodendritic damage, including the loss of dendritic spines in brain regions critical for learning and memory (40–43). Furthermore, plasma biomarkers point to underlying immune and metabolic dysregulation. For example, altered levels of triggering receptor expressed on myeloid cells 2 (TREM2), a macrophage receptor dysregulated in the brains of PWH with NCI, and growth differentiation factor 15 (GDF15), a mitokine associated with metabolic stress, implicate these pathways in NCI pathogenesis (44). Given that cannabinoids can promote immune and metabolic homeostasis, changes in TREM2 and GDF15 levels in cannabis-using PWH may reflect better cellular and metabolic balance. In parallel, plasma biomarkers of neuronal health, such as brain-derived neurotrophic factor (BDNF), directly correlate with neurocognitive function (45–47). Therefore, the plasma profile of PWH can serve as a valuable indicator of their immune dysregulation and neuronal health.

This exploratory study investigated the impact of cannabis use on MDMs and neurocognition in PWH. The primary objective was to characterize cannabis-related immunometabolic changes in MDMs by examining mitochondrial function and gene expression across different groups (PWoH, PWH, and frequency of cannabis use (naïve/low, moderate, daily)). The secondary aims assessed whether cannabis use is associated with lower NCI in PWH as well as quantifying related plasma biomarkers. As such, this work offers clinical relevance by potentially revealing a mechanistic link that mediates the neuroprotective and disease-modifying effects of cannabis in PWH. The findings could, therefore, support the hypothesis that cannabis use mitigates NCI in PWH by reprogramming MDM function toward an anti-inflammatory and neuroprotective phenotype.

## Materials and Methods

### Sex as a Biological Variable

This study included both male and female humans, and similar findings are reported for both sexes.

### Study Population

**Graphical Methods**

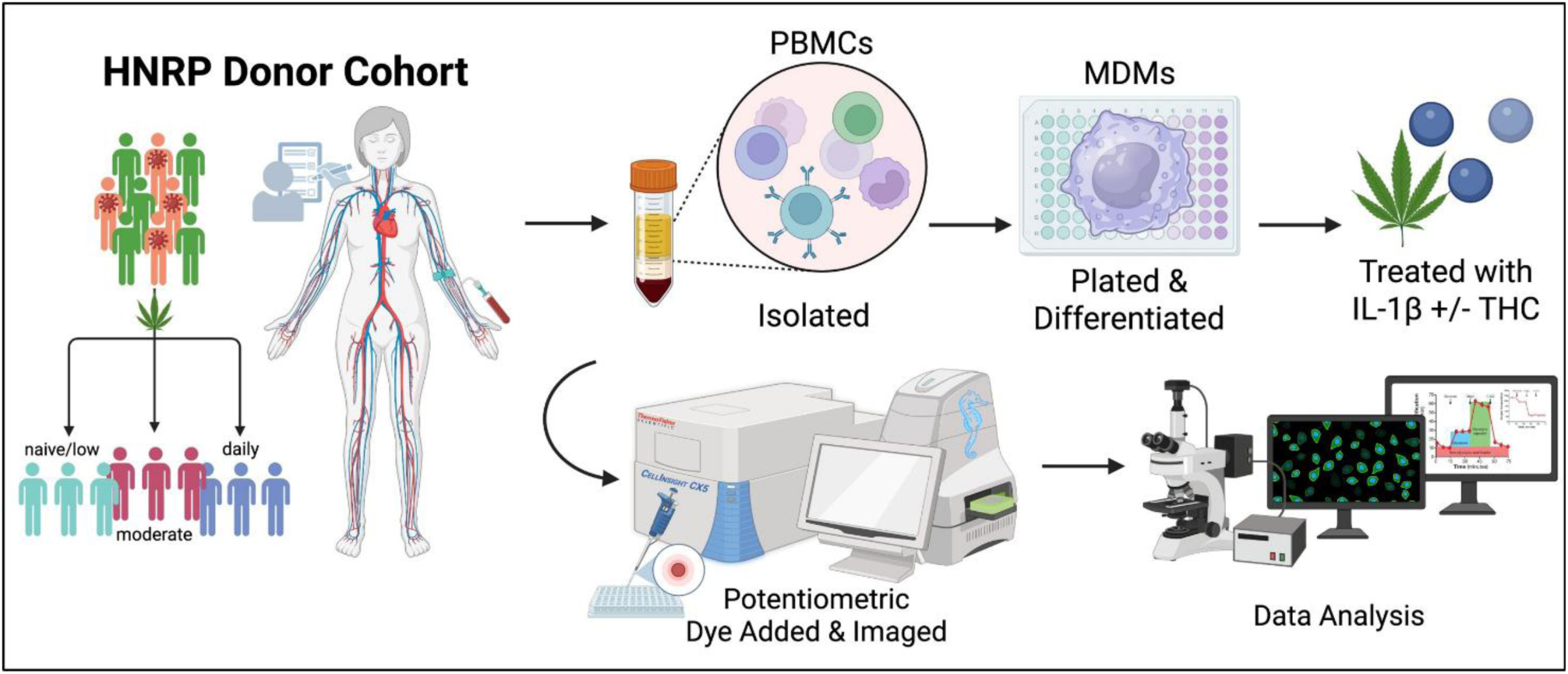

From 2023 to 2025, this study recruited individuals with and without HIV from San Diego, CA, and categorized them according to frequency of cannabis use in the six months prior to their clinical assessment visit at the HIV Neurobehavioral Research Program (HNRP). Categorization was as follows: Naïve/Low (never used or used ≤ 6 times/year with > 60 days abstinence), Moderate (1–6 times/week), and Daily (7 days/week). Current and lifetime cannabis use was assessed using both laboratory measures and self-report questionnaires to capture variations in frequency, quantity, and cannabinoid content. To control for potential withdrawal effects, cannabis users were instructed to maintain their usual consumption before assessment.

Participants provided venous blood samples in addition to performing comprehensive medical and neurobehavioral evaluations at the HNRP. Exclusion criteria included acute substance use (non-cannabis) as indicated by urine drug screen and breathalyzer on the day of assessment, significant or uncontrolled medical/psychiatric/neurological conditions, use of daily anti-inflammatory medication, DSM IV diagnosis of significant (i.e., dependence) substance use disorders within the past five years, DSM diagnosis of moderate or severe alcohol use disorder in the past 12 months, or mild (i.e., abuse) substance use disorder within the past six months. Substance-related criteria apply to substances other than cannabis for the cannabis-using groups and tobacco. A total of 83 eligible participants were included that were virally suppressed: PWH cannabis users (n = 27), PWH cannabis-naïve (n = 24), PWoH cannabis users (n = 18), and PWoH cannabis-naïve (n = 15) (**Table 1**). Among cannabis users in the cohort, nearly half (46%) reported using more than one route of administration, reflecting common real-world use patterns. Among the remaining participants who reported a single primary route, smoking was most common (46%), followed by ingestion (32%) and vaping (22%). Importantly, the distribution of routes of administration did not differ significantly between daily and moderate cannabis users (*p*>0.10).

**Table 1.**
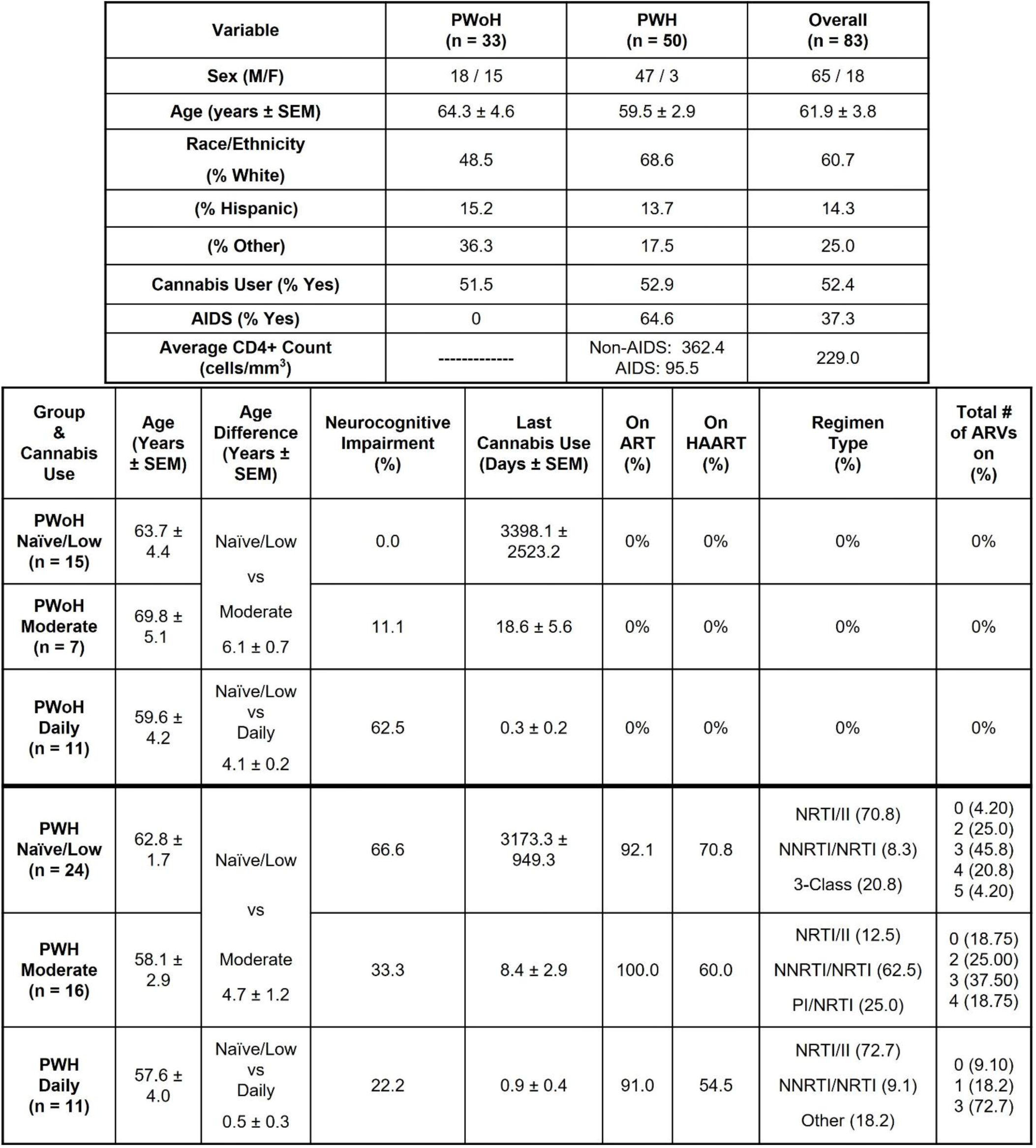
Clinical data of sampled patients showing the Sex, Age, and Ethnicity of the PWoH, PWH, and total overall groups. The table is further organized for Age, Age difference, Neurocognitive Impairment (%), Last cannabis use, on ART (%), on HAART (%), Regimen type (%), Total # of ARVs on (%).

### Separation and Treatment of Monocyte-Derived Macrophages (MDMs)

Peripheral blood mononuclear cells (PBMCs) were isolated from participant blood samples using density gradient centrifugation. Two hours after collection, 45 mL of donor blood was gently layered into three 50 mL conical tubes containing HISTOPAQUE-1077 (Sigma Life Sciences; Cat. # 10771) at a 1:1 ratio. This was followed by centrifugation at 400g for 30 minutes with no brake. After centrifugation, the PBMC layer was collected, diluted 1:1 with 1X Phosphate Buffered Saline (PBS), and centrifuged at 250g for 10 minutes. The cells were washed, centrifuged, and resuspended three times in 1X PBS before being resuspended in Iscove’s Modified Dulbecco’s Medium (IMDM; Gibco; Cat. # 12440053) supplemented with 10% human serum (Millipore Sigma; Cat. # H5667) and 1% penicillin/streptomycin (Gibco; Cat. # 15140122). Cell counts were performed using a Countess™ 3 FL automated cell counter (Thermo Fisher Scientific; Cat. # AMQAF2000) with 0.4% trypan blue solution (Amresco; Cat. # K940100ML). The cells were plated in 96-well plates (Thermo Fisher Scientific; Cat. # 164588) at a density of 1 × 10⁴ cells/well for immunocytochemistry analysis. MDMs from donors representative of each demographic group were also plated in Seahorse XFe96/XF Pro Cell Culture Microplates (Agilent Technologies; Cat. # 103794-100) at a density of 3 × 10⁴ cells/well for Seahorse analysis. The cells were cultured and differentiated in a humidified incubator at 5% CO2 and 37°C. Monocytes were isolated via plastic adhesion and matured into monocyte-derived macrophages (MDMs) in supplemented IMDM media. Non-adherent cells were removed through complete media exchanges every two to three days. After eight days, MDMs were pre-treated for one hour with delta-9-tetrahydrocannabinol (THC, Millipore-Sigma), before being incubated with IL-1β (InvivoGen; Cat. # 6409-44-01; 20 ng/mL) for 24 hours. All conditions were tested in biological replicates of at least 6.

#### Mitochondrial Signal Measurements

Mitochondria signal was measured using cells preincubated with MitoTracker™ Deep Red FM (MT, Invitrogen, Cat. # M22426), following the manufacturer’s instructions for immunocytochemistry. After 24 hours of treatment, MDMs were washed with 1x PBS. Then the cells were incubated with 50 µL of 100 nM MT at 37°C with 5% CO2. After 30 minutes, wells were washed three times with 1x PBS before being fixed in 50 µLs of 4% paraformaldehyde for 20 minutes at 4°C. Prior to imaging, the cells were incubated on a shaking platform for 5 minutes at room temperature with 4′,6-diamidino-2-phenylindole (DAPI) to localize cellular nuclei then washed with 1x PBS three times. Cells were visualized and imaged using two channels on the CellInsight CX5 high-content screening platform (Thermo Fisher Scientific, Cat. # CX51110).

DAPI was imaged on Channel 1 at 358nm excitation and 461nm emission and used to autofocus the CX5 machine to the plane of interest at a using a 20X objective lens. MT was imaged on Channel 2 using 633 nm excitation and 650-734 nm emission. Each well was imaged twice across the two channels with five different fields of view per well. Image exposure time was optimized to each plate using a uniform threshold intensity and a 40% fluorescence capture target.

The Thermo Scientific HCS Studio 4.0 Cell Analysis Software, included with the CX5 platform, was used to configure assay parameters as well as tabulate results and population statistics for all wells imaged. Representative images from each plate were used to validate and optimize two assays: the Cell Health Profiling Assay and the Spot Detection Assay. Images were preprocessed on both Channel 1 (DAPI) and Channel 2 (MT) to eliminate any background signal detected in control wells. In Channel 1, primary objects were identified using thresholding and segmentation. Objects were validated by gating for area, average intensity, and variable intensity. Channel 2 objects were selected by adjusting a ring mask around validated Channel 1 objects, then gating the mask for upper and lower parameters of object total intensity and average intensity measurements. Spots within each Channel 2 ring were then identified using Box Detection, Three Sigma Thresholding, and Uniform Smoothing. The spots were then gated for upper and lower parameters using area, shape, average intensity, and variable intensity. After assay parameters were configured, two scans of each plate were conducted.

For Spot Average Intensity (the average individual mitochondrial intensity per well), Spot Average Area (the average individual mitochondrial size per well), Spot Total Area per Object (the average mitochondrial area per cell), and Target Average Intensity (the average MT intensity per cell, all values were measured in pixels.

In order to normalize values between plate reads, the pixel values for each plates’ control wells were averaged. All well values scanned were then divided by the average pixel value of the control wells, for all image features. The values for each treatment were then averaged for group comparisons. Calculations were performed using Excel and PRISM, with the ROUT method used to identify and exclude outliers.

#### Determination of Metabolism in Monocyte-Derived Macrophages

Max ECAR and Max OCAR output from MDMs was determined using the Seahorse XF Pro Analyzer (Agilent Technologies, Santa Clara, CA), courtesy of the Mitochondrial Bioenergetics Resource Core at the University of California, San Diego School of Medicine (La Jolla, CA). The Mitochondrial Bioenergetics core kindly provided the sensor cartridges, tissue culture plates, calibrant, and utility plates for calibration. Seahorse data was normalized using direct cell count, either via Agilent’s XF Imaging system or post-assay fluorescent DAPI staining. Normalization was performed using the Seahorse Wave Desktop software (Agilent Technologies, Santa Clara, CA). The Seahorse Wave calculated the Max OCR and Max ECAR values for each well, and the values for which were then exported into PRISM. Outliers were removed via the PRISM ROUT method before well values were averaged per treatment. Perimeter well values were excluded from analysis.

#### RNA Sequencing

RNA sequencing on MDMs of all demographic groups was conducted at the Institute for Genomic Medicine, University of California, San Diego (La Jolla, CA). An Agilent Tapestation (Santa Clara, CA) automated gel electrophoresis machine was used for sample quality control before samples were sequenced using an Illumina NovaSeq X Plus.

After sequencing, biosample fastq files were uploaded to the BaseSpace Sequence Hub (Illumina, San Diego, CA). The BaseSpace STAR aligner (v2.7.9a) was used to align clean reads to the hg38 human reference genome with default parameters.

Differential expression analyses were run between all sample groups using the BaseSpace DRAGEN Differential Expression Application (v4.3.7). See Statistical Analysis (in Methods) for further bioinformatic analysis details.

#### Quantitative Polymerase Chain Reaction (qPCR)

RNA isolation and cDNA synthesis on MDMs representative of each demographic group was performed as previously described [37]. Multiplex relative quantification assays were performing using a QuantStudio 3 Real-Time PCR machine using TaqMan Fast Advanced Master Mix for qPCR (Applied Biosystems; #4444557). ACTB (Applied Biosystems; #4310881E) was used as the endogenous control. Individual probes of interest included MT-CO3 (Life Technologies, Cat.# 448892), PGC-1α (Life Technologies, Cat.# 4331182), PINK1 (Life Technologies, Cat.# 4331182), PGAM1 (Fischer Scientific, Cat.# 4351372), TPI1 (Life Technologies, Cat.# 4351372), GPI (Life Technologies, Cat.# 4351372), MT-CO1 (Life Technologies, Cat.# 4331182), MT-CYB (Life Technologies, Cat.# 4331182), BDNF (Life Technologies, Cat.# Mm04230607_s1), IL-6 (Life Technologies, Cat.# 4414149), IL-1α (Life Technologies, Cat.# Mm00439620_m1), IL-18 (Life Technologies, Cat.# Mm00434226_m1), CD68 (Life Technologies, Cat.# Mm00839636_g1), IL-1β (Life Technologies, Cat.# Hs01555410_m1), IL-4 (Life Technologies, Cat.# 4414148), IL-10 (Life Technologies, Cat.# 4414143), and CD163 (Life Technologies, Cat.# Hs00174705_m1). Raw cq data was uploaded to the ThermoFisher Connect cloud before being exported for analysis.

The gene expression fold change was quantified via the comparative Ct method, as previously described with outliers removing using the ROUT method [64]. All donor samples were run in technical triplicate.

#### Differentially Expressed Genes Displayed

Gene analyses as in **Figure 2D–2G** display significant differentially expressed genes (DEGs) identified across group comparisons, such as PWH Naïve vs. PWoH Naïve. Consistent parameters were applied across all panels; the genes meeting the DEG criteria for a specific category (e.g., Mitochondrial Biogenesis) were included in the respective analysis and also including non-DEGs to demonstrate a robust list.

**Figure 1.**
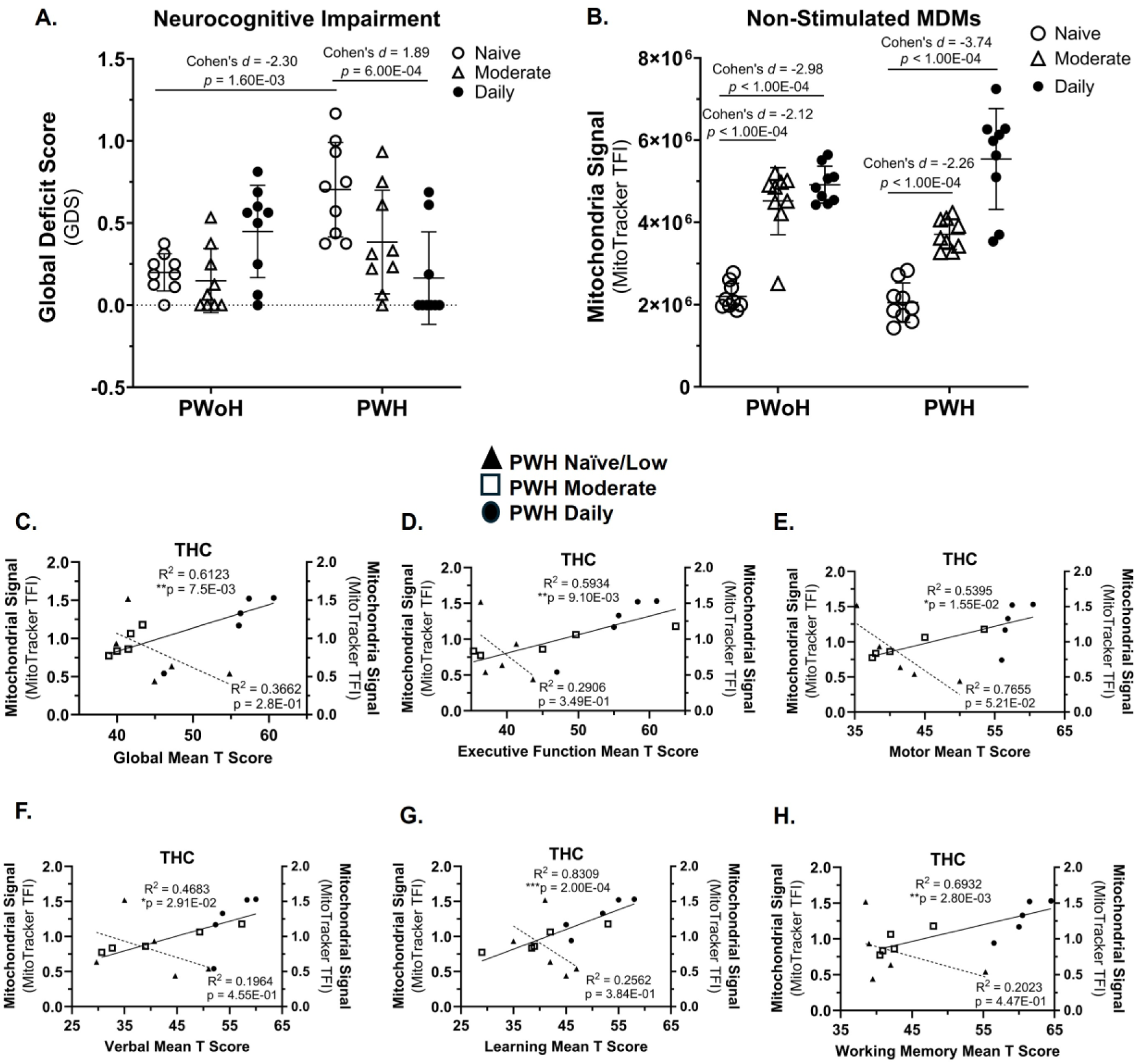
Cannabis use by PWH was associated with better neurocognitive function that correlated with higher mitochondrial signals in MDMs. (**A**) NCI was clinically assessed using the Global Deficit Score (GDS, 0 = no impairment, 1.5 = severe impairment). (**B**) Mitochondria signal in non-stimulated MDMs was quantified via MitoTracker total fluorescence intensity (TFI) on the CX5. (**C–H**) Correlations with THC-stimulated MDMs included GDS, Executive Function, Motor skills, Verbal skills, Learning, and Working Memory. *Statistics*: Two-way ANOVA with Tukey’s Multiple Comparisons Test. Simple Linear Regression Analysis. (B) Mitochondrial Signal: Inter-assay CV = 9.2%; Mitochondrial Signal LLOQ: 10 nM; GDS PWoH Naïve/Low vs PWH Naïve/Low: Cohen’s *d* = -2.30 and effect-size *r* = -0.76; GDS PWH Naïve/Low vs PWH Daily: Cohen’s *d* = 1.89 and effect-size *r* = 0.69; Mitochondrial Signal PWoH Naïve/Low vs PWH Naïve/Low: Cohen’s *d* = -6.94 and effect-size *r* = -0.96; Mitochondrial Signal PWH Naïve/Low vs PWH Daily: Cohen’s *d* = -3.74 and effect-size *r* = -0.88. GDS PWoH Naïve/Low vs PWH Naïve/Low: 95% CI [-0.71, -0.3]; GDS PWH Naïve/Low vs PWH Daily: 95% CI [0.27, 0.8]; Mitochondrial Signal PWoH Naïve/Low vs PWH Naïve/Low: 95% CI [-3074198.2, -2352554.9]; Mitochondrial Signal PWH Naïve/Low vs PWH Daily: 95% CI [-4354553.22, -2629851.48]; Global Mean T Score: 95% CI [0.01078, 0.05059], effect-size *r* = 0.97; Exec. Function Score: 95% CI [0.2732, 0.9427], effect-size *r* = 0.77; Motor Score: 95% CI [0.1951, 0.9328], effect-size *r* = 0.73; Verbal Score: 95% CI [0.09608, 0.9183], effect-size *r* = 0.68; Learning Score: 95% CI [0.6617, 0.9792], effect-size *r* = 0.91; Working Memory Score: 95% CI [0.4266, 0.9593], effect-size *r* = 0.83; (A-B) n = 9 donors per group; (C – H) n = 5 donors per group.

**Figure 2.**
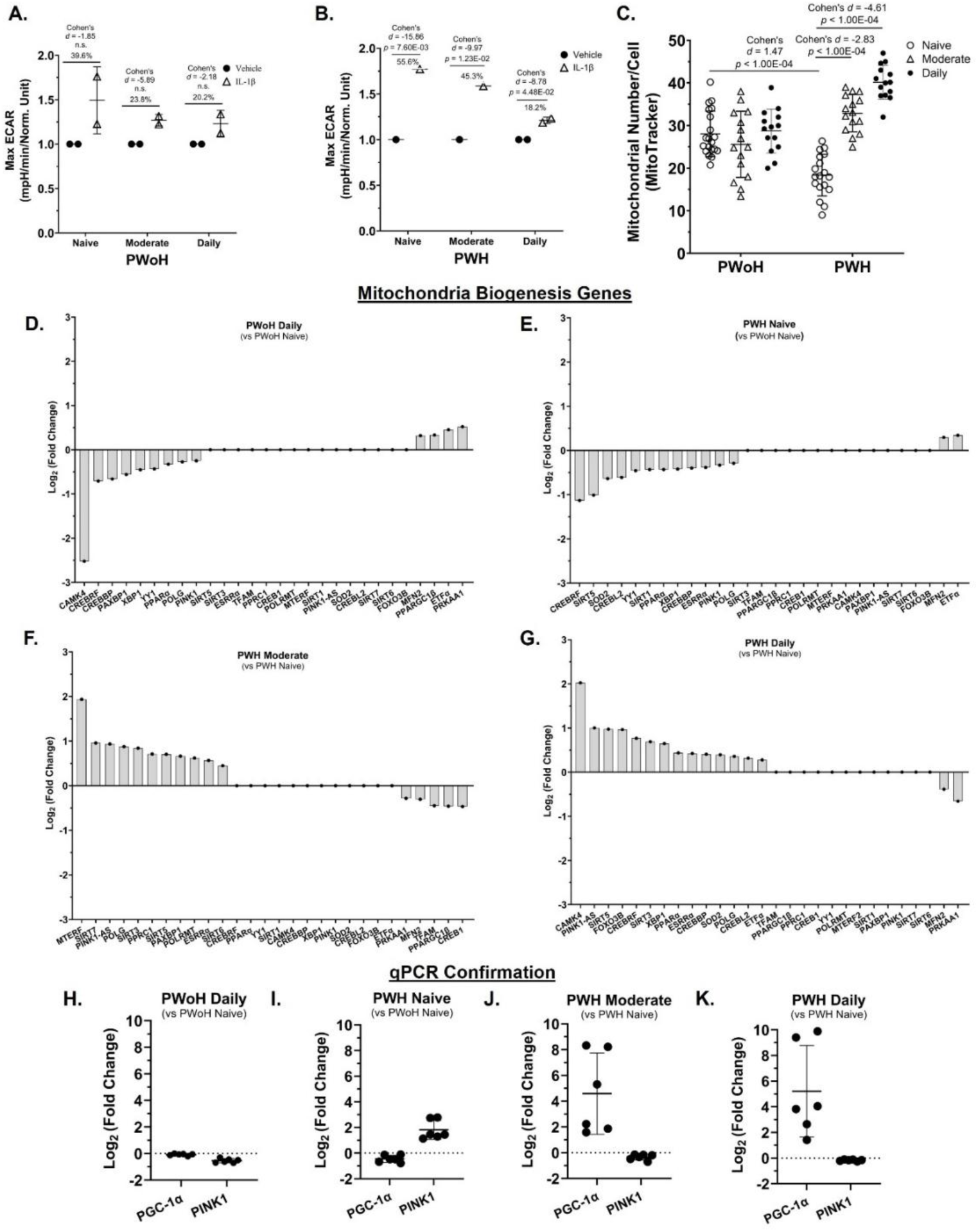
Cannabis use by PWH was associated with a reduced glycolytic response, increased mitochondrial numbers and transcriptional profiles consistent with mitochondrial biogenesis in MDMs. (**A** and **B**) Glycolytic responses were measured via a Seahorse assay (Max ECAR) following IL-1β stimulation. (**C**) Mitochondrial numbers per MDM was quantified via MitoTracker staining and CX5 imaging. (**D–G**) Mitochondrial biogenesis gene expression was determined using RNA sequencing. (**H–K**) Gene expression for PGC-1α and PINK1 was confirmed via qPCR (Log2 Fold Change). ECAR: Extracellular Acidification Rate. *Statistics*: Two-way ANOVA with Tukey’s Multiple Comparisons Test. Max ECAR: Intra-assay CV = 5.1% and 2.8%, Inter-assay CV = 4.0%; Mitochondrial Number/Cell: Inter-assay CV = 7.9%; (H – K) qPCR: Inter-assay CV = 6.4%; RNA sequencing LLOQ: 1 pg; qPCR LLOQ: 10 copies of target DNA; CX5 Mitochondrial Number LLOQ: 10 nM; Max ECAR PWoH Daily Control vs IL-1β: 95% CI [-0.5683, -0.09420], effect-size *r* = 0.74; Max ECAR PWH Daily Control vs IL-1β: 95% CI [-0.5998, -0.4461], effect-size *r* = 0.98; Mitochondrial Numbers PWoH Naïve/Low vs PWH Naïve/Low: 95% CI [4.04, 12.28], effect-size *d* = 1.47, *r* = 0.59; Mitochondrial Numbers PWH Naïve/Low vs PWH Daily: 95% CI [-25.56, -18.48], effect-size *d* = -4.61, *r* = -0.92; PGC-1α PWoH Daily vs PWH Naïve/Low: 95% CI [-0.61, -0.11], effect-size *d* = -1.78, *r* = -0.67; PGC-1α PWoH Daily vs PWH Moderate: 95% CI [-7.27, -1.74], effect-size *d* = - 2.02, *r* = -0.71; PGC-1α PWoH Daily vs PWH Daily: 95% CI [-8.25, -1.99], effect-size *d* = -2.03, *r* = -0.71; (A and B) n = 2 donors per group (PWoH and PWH: Naïve/Low, Moderate, Daily); (D -G) n = 1 donor per group with 3 biological replicates; (H – K) n = 3 donors with 2 biological replicates per group.

For instance, *CREBRF* met the threshold for differential expression in panels **2D**, **2E**, and **2G**, but did not reach significance in **2F** but was now included in **2F**. To illustrate the directionality of these changes, the DEGs in panels **2D** and **2E** are ordered from lowest to highest expression (downregulated), while panels **2F** and **2G** are ordered from highest to lowest (upregulated).

In **Figures 3J – 3M** and **4I – 4L**, while most genes remain consistent, the panels specifically highlight significant DEGs to illustrate metabolic shifts. **Figure 3** (Glycolysis): Genes in panels **3J – 3K** are ordered from highest to lowest expression to demonstrate the upregulation of glycolysis, whereas panels **3L – 3M** are ordered from lowest to highest to highlight the downregulation of glycolytic genes. **Figure 4** (Oxidative Stress): Significant DEGs in panels **4I – 4P** are organized from highest to lowest expression. This arrangement demonstrates that oxidative stress genes are upregulated in PWoH Daily and in PWH Naïve groups. Whereas these genes become downregulated in PWH who are moderate and daily cannabis users. In both figures, the data is structured to show how significant DEGs related to oxidative stress and glycolysis in PWH are attenuated with cannabis use. Because we prioritized displaying the significant DEGs for each comparison, the specific gene lists vary slightly between categories, though the majority of genes are consistent across the analyses.

**Figure 3.**
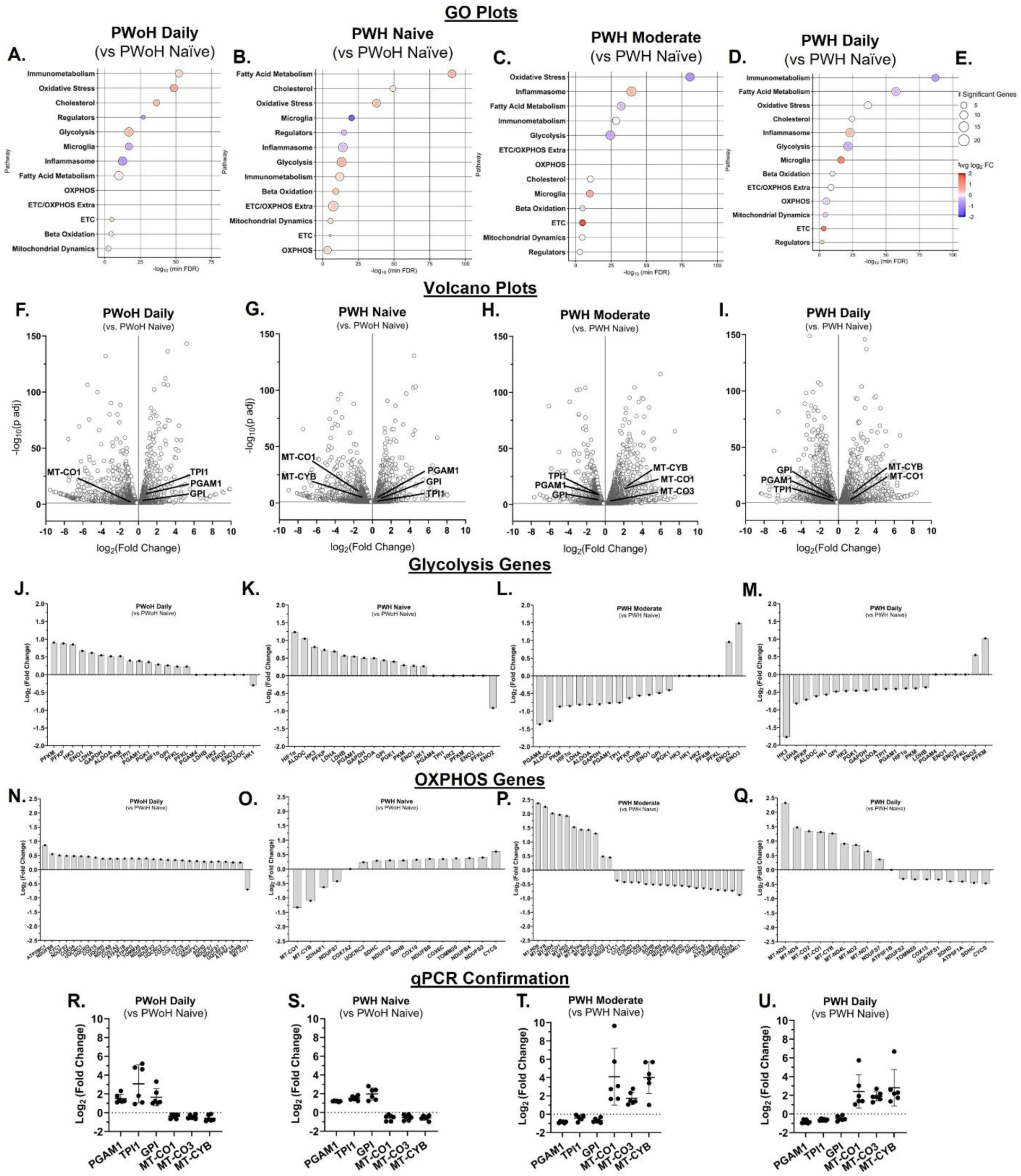
Cannabis use by PWH was associated with lower glycolysis and higher oxidative phosphorylation gene expression in MDMs. (**A–E**) Gene Ontology (GO) enrichment analysis of metabolic pathways. (**F–I**) Volcano plot visualization of upregulated and downregulated gene expression including glycolysis and oxidative phosphorylation (OXPHOS) gene expression. (**J–Q**) RNA sequencing determination of glycolytic and OXPHOS gene expression. (**R–U**) Confirmation of randomly selected three glycolysis and three OXPHOS genes via qPCR (Log2 Fold Change). (R-U) qPCR: Inter-assay CV = 4.1%; RNA sequencing LLOQ: 1 pg; qPCR LLOQ: 10 copies of target DNA; GO: Gene Ontology. (A – Q) n = 1 donor per group with 3 biological replicates (PWoH and PWH: Naïve/Low, Moderate, Daily); (R – U) n = 3 donors with 2 biological replicates per group.

**Figure 4.**
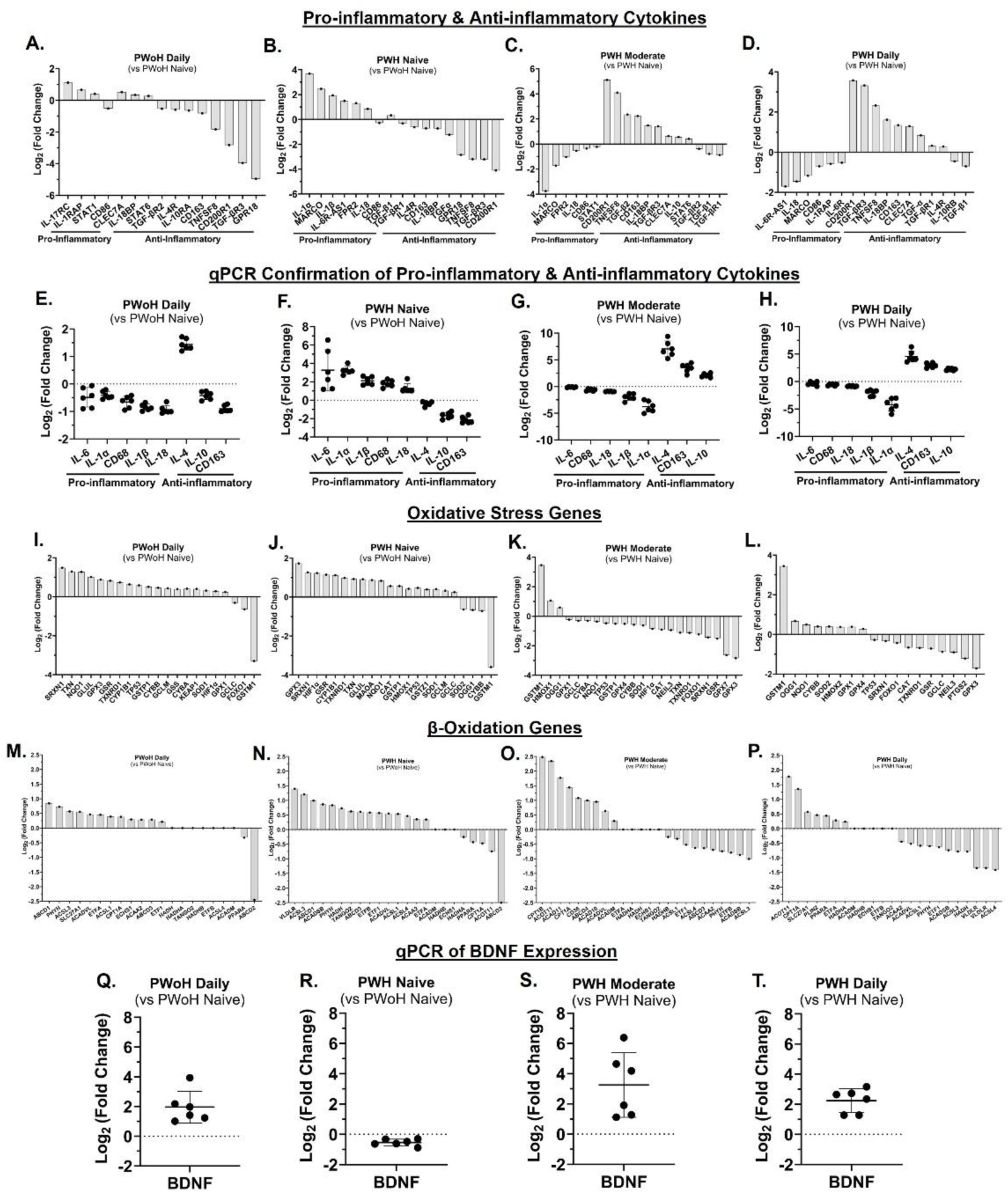
Cannabis use by PWH was associated with reduced pro-inflammatory and oxidative stress gene expression and increased anti-inflammatory, β-oxidation, and brain-derived neurotropic factor gene expression. (**A–H**) Gene expression analysis (RNA sequencing and qPCR) of pro-inflammatory and anti-inflammatory cytokine expression in MDMs from PLWoH and PWH (cannabis-naïve, moderate, and daily users). (**I–L**) RNA sequencing analysis of oxidative stress gene expression. (**M–P**) RNA sequencing analysis of β-oxidation gene expression. (**Q–T**) qPCR analysis of BDNF gene expression (Inter-assay CV = 5.9%). BDNF: Brain-Derived Neurotrophic Factor; LLOQ: Lower Limit of Quantitation; RNA sequencing LLOQ: 1 pg; qPCR LLOQ: 10 copies of target DNA; *Statistics*: BDNF PWoH Daily vs PWH Naïve/Low: 95% CI [1.55, 3.45], effect-size *d* = 3.25, *r* = 0.85; BDNF PWoH Daily vs PWH Moderate: 95% CI [-3.39, 0.8], effect-size *d* = -0.77, *r* = -0.36; BDNF PWoH Daily vs PWH Daily: 95% CI [-1.44, 0.88], effect-size *d* = -0.30, *r* = -0.15; (A – D, I – P) n = 1 donor per group with 3 biological replicates (PWoH and PWH: Naïve/Low, Moderate, Daily); (E – H) n = 3 donors with 2 biological replicates per group; (Q – T) n = 6 donors per group.

#### Clinical Assessment

Clinical data were obtained through standardized neuro-medical, neurocognitive, and psychiatric/substance use assessments.

*Neuromedical Assessment.* A structured medical history interview was conducted to evaluate both current and past medical conditions (e.g., Hepatitis C Virus [HCV], diabetes, hypertension, hyperlipidemia) and treatment. Antiretroviral (ARV) treatment history was assessed using a structured, clinician-administered questionnaire.

Participants also completed a breathalyzer test to detect recent alcohol use, and blood and urine samples were collected for routine clinical labs (e.g., lipid panel, CD4^+^T-cell counts, plasma HIV RNA, C Reactive Protein) and research biomarkers (see under: Plasma Biomarkers), diagnostic testing (e.g., HIV, HCV), and urine toxicology screening. None of the participants had a positive breathalyzer result on the morning of their evaluation. Health-related quality of life was assessed using the 36-Item Short Form Health Survey (SF-36) (48), which measures eight domains of functioning and well-being. Scores were calculated according to standardized procedures, with higher values indicating better health-related quality of life.

#### Detecting Plasma Biomarkers

Enzyme-Linked Immunosorbent Assay (ELISA) kits were used to measure plasma levels of Human GDF15 (Thomas Scientific, Cat.# KE00108-96T), Human soluble TREM2 (Invitrogen, Cat.# EH464RB), Human Mature BDNF (Biosensis, Cat.# BEK-2211-1P), and Human proBDNF (Biosensis, Cat.# BEK-2237-1P) according to the manufacturer’s instructions, and normalized to standard curves. Fluorescence of the product of the enzyme reaction was detected using a plate reader (Beckman Coulter DTX 880 multimode microplate reader, US).

#### Neurocognitive Assessment

Participants completed a standardized battery of 14 tests designed to assess seven cognitive domains relevant to HIV and/or cannabis use (30,49): verbal fluency, information-processing speed, executive functions, learning, memory, working memory, and motor skills. Raw scores from each test were converted to demographically adjusted T-scores (Mean=50; SD=10) based on published normative samples that account for age, education, sex, and race/ethnicity (50–52). Domain-specific T-scores were calculated by averaging individual test T-scores within each domain, and a global neurocognitive T-score was derived by averaging across all tests. Higher T-scores indicated better neurocognitive performance. Demographically corrected T-scores were converted to deficit scores ranging from 0 (T > 39, no impairment) to 5 (T < 20, severe impairment). These were averaged to yield domain deficit scores (DDS) and a global deficit score (GDS), which served as outcome measures in analyses. Global neurocognitive impairment was classified using a GDS cutoff of ≥0.5, representing at least mild impairment on half of the battery, and domain impairment was defined by a DDS cutoff of >0.5 (53).

#### Psychiatric and Substance Use History

Current and lifetime substance use (i.e., DSM-IV Substance Abuse/Dependence) and mood disorders were assessed using the Composite International Diagnostic Interview (54). Lifetime substance use characteristics (e.g., age at first and last use, frequency, quantity) were collected using a semi-structured Timeline Follow-Back substance use interview.

#### Statistical Analysis

Statistical analyses were performed using Microsoft Excel, GraphPad Prism 10 (La Jolla, CA), as well as R (4.4.3) and RStudio (2025.05.1+513).

##### Group Comparisons

Differences in in vitro treatment effects and plasma analyte levels across demographic groups were assessed using two-sample t-tests, one-way ANOVA, or two-way ANOVA followed by Tukey’s post hoc pairwise comparisons, unless otherwise specified. Two-tailed t-tests were applied for pairwise comparisons involving MT signal, treatment conditions, and demographic variables. Experimental sample sizes and details regarding data normalization to appropriate controls are provided in the corresponding figure legends. Statistical significance was set at p < 0.05.

##### Correlational Analysis

Only participants with complete clinical datasets and corresponding MT measurements were included in the analyses. Simple linear regression was employed to examine the relationships between MT signal measures and clinical variables (e.g. GDS). The regression line range was automatically determined by PRISM with the 95% confidence bands visualizing the line of best fit. Effect sizes were reported using R² values.

##### Data Normalization and Visualization

MT signal measures were normalized to control MDMs treated with media. A correlation matrix was generated to visually represent the relationships between MT and clinical data. These models controlled for the influence of key covariates on the observed associations.

##### Gene Ontology and Volcano Plots

Plots of the DRAGEN Differential Expression datasets were generated using RStudio, Excel and Prism. For the GO Plots, CSV files of differentially expressed genes (DEGs) with False Discovery Rate-adjusted p-values were downloaded from BaseSpace and read into the R data frame. Genes displaying a false discovery rate (FDR) < 0.05 were considered significantly differentially expressed. Analyses were conducted using open-source packages on R, including clusterProfiler, org.Hs.eg.db, dplyr, ggplot2, and enrichplot with target genes of interest determined by the authors. Gene ontology dot plots were generated primarily using the ggplot2 package in R Studio.

For the Volcano Plots, datasets were downloaded from BaseSpace, and genes with missing adjusted p-values (NaN) or adjusted p-values greater than 0.05 were excluded from further analysis. No additional filtering was applied based on log₂ fold change.

Adjusted p-values (p-adj) were transformed to –log₁₀(p-adj) values in Microsoft Excel, and volcano plots were generated using PRISM. Genes of interest highlighted in the plots were selected from a researcher-curated list.

The Log2 Fold Change plots on Glycolysis genes, OXPHOS genes, Pro- & Anti-Inflammatory Cytokines, Oxidative Stress genes and B-Oxidation genes were created using Log₂ Fold Change data obtained directly from BaseSpace, restricted to a researcher-curated list.

## Results

### Cannabis use by PWH was associated with better neurocognitive function that correlated with higher average mitochondrial signals in monocyte-derived macrophages (*Figure 1*)

The relationship between HIV status and the Global Deficit Score (GDS), is significantly moderated by cannabis use frequency, as indicated by a significant interaction effect (*F*(2,48) = 11.00, *P* = 0.0001).

Post hoc analyses showed that cannabis-naïve/low PWH demonstrated significantly greater global NC deficits than cannabis-naïve/low PWoH (*p* = 1.60 × 10⁻^3^, 95% CI [−0.71, −0.3]). Among PWH, daily cannabis users had significantly fewer global NC deficits than cannabis-naïve/low PWH (*p* = 6.00 × 10⁻^4^, 95% CI [0.27, 0.8]) (Figure 1A). A significant interaction was observed between HIV status and cannabis use frequency concerning mitochondrial signal levels (*F*(2,48) = 4.870, (*P* = 0.0119). This interaction indicates that the effect of HIV status on mitochondrial signal depends on the frequency of cannabis use (Figure 1B).

Post-hoc analysis showed that within both PWoH and PWH groups, moderate and daily cannabis users had a significantly higher mitochondrial signal compared to cannabis-naïve/low users (*p* < 1.00 × 10^-4^), 95% CI [-4,354,553.22, -2,629,851.48]) (Figure 1B).

To examine the relationship between mitochondrial function following THC exposure and neurocognitive performance in cannabis-using PWH, MDMs were incubated with THC, MitoTracker signal was quantified, and mitochondrial signal was correlated with global and domain-specific T-scores. This exposure resulted in a positive correlation between mitochondrial signal and improved cognitive performance across several functions. Significant correlations were found for Global Mean T Score (*p* = 7.50 × 10^-3^), 95% CI [0.01078, 0.05059]) and domain T-scores, including Executive Function (*p* = 9.10 × 10^-3^, 95% CI [0.2732, 0.9427]), Motor Skills (*p* = 1.55 × 10^-2^, 95% CI [0.1951, 0.9328]), Verbal Ability (*p* = 2.91 × 10^-2^, 95% CI [0.09608, 0.9183]), Learning (*p* = 2.00 × 10^-4^, 95% CI [0.6617, 0.9792]), and Working Memory (*p* = 2.80 × 10^-3^, 95% CI [0.4266, 0.9593]) (Figure 1, C–H). No significant relationship was detected between MDM Mitochondrial signal and any cognitive score within the PWH naïve group (Figure 1, C–H). These results indicate a potential link between daily cannabis use, better neurocognition, and a higher mitochondrial signal in MDMs from PWH.

### Cannabis use by PWH was associated with a reduced glycolytic response, increased mitochondrial numbers and transcriptional profiles consistent with mitochondrial biogenesis in monocyte-derived macrophages (Figure 2)

A significant interaction was observed between PWH and frequency of cannabis use with Max ECAR for IL-1β treatment in MDMs (*F*(2, 2) = 103.1, *P* = 0.0096), but there was no significant interaction between PWoH and frequency of cannabis use with Max ECAR for IL-1β treatment in MDMs (*F*(2, 6) = 0.7197, *P* = 0.5246). This finding indicates that the effect of PWH on Max ECAR is dependent on the frequency of cannabis use (Figure 2A).

In MDMs from PWH, daily cannabis was associated with significantly diminished IL-1β-stimulated glycolysis compared to naïve/low (*p* = 4.48 × 10^-2^, 95% CI [-0.5998, - 0.4461]). This relationship was not observed in MDMs from PWoH (Figure 2, A and B). Consistent with these metabolic changes, mitochondrial numbers per MDM were significantly higher in moderate and daily cannabis-using PWH relative to cannabis-naïve/low PWH (*p* < 1.00 × 10^-4^, 95% CI [-25.56, -18.48]; Figure 2, C).

Gene expression analyses supported these findings. RNA sequencing showed that mitochondrial biogenesis genes that showed lower expression in cannabis-naïve/low PWH compared to cannabis-naïve/low PWoH (Figure 2, D & E) showed higher expression in moderate or daily-using PWH compared to their cannabis-naïve/low PWH counterparts (Figure 2, F & G).

Further supporting this, qPCR analysis confirmed a significantly higher mRNA levels of the key mitochondrial biogenesis regulator, PGC-1α, in moderate and daily cannabis-using PWH (up to 8 and 10 Log2 fold change, respectively) when compared to both daily cannabis-using PWoH and cannabis-naïve/low PWH (Figure 2, H – K).

Additionally, PINK1 mRNA, coding for a gene that detects damaged mitochondria, was higher in cannabis-naïve/low PWH relative to PWoH but was lower in moderate and daily cannabis-using PWH (Figure 2, I – K).

Thus, these findings indicate that moderate and daily cannabis use by PWH is associated with reduced levels of mRNA associated with glycolysis and higher mitochondrial numbers, a change supported by the upregulation of PGC-1α and a positive correlation with improved neurocognitive outcomes.

### Cannabis use by PWH was associated with lower glycolysis and higher oxidative phosphorylation gene expression in monocyte-derived macrophages (Figure 3)

Gene expression analysis of MDMs revealed a metabolic shift influenced by both cannabis use and HIV status. In daily cannabis-using people without HIV (PWoH) and cannabis-naïve/low people with HIV (PWH), Gene Ontology (GO) and volcano plots showed a significant enrichment of glycolysis-related genes compared to cannabis-naïve/low PWoH (Figure 3, A, B, F, and G). Genes associated with oxidative phosphorylation (OXPHOS), particularly the electron transport chain (ETC), were not significantly enriched. This higher level of glycolytic gene expression was consistent with transcripts identified by RNA sequencing, which also showed higher expression of OXPHOS genes (Figure 3, J–Q). These transcript profiles were validated by qPCR analysis showing higher expression of glycolytic genes (*PGAM1, TPI1, GPI*) and lower expression of OXPHOS genes (*MT-CO1, MT-CO3, MT-CYB*) (Figure 3, R and S).

Conversely, MDMs from PWH with moderate or daily cannabis use exhibited a different metabolic profile. Compared to cannabis-naïve/low PWH, these cells showed significant enrichment of OXPHOS and ETC genes, with glycolysis genes not being enriched (Figure 3, C, D, H, and I). RNA sequencing confirmed this metabolic reversal (Figure 3, J–Q), and qPCR validated the transcript profile, showing lower glycolytic gene expression and higher OXPHOS gene expression (Figure 3, T and U).

These findings collectively demonstrate that cannabis use by PWH is associated with increased mitochondrial oxidative phosphorylation in MDMs, suggesting a metabolic reprogramming away from glycolysis.

### Cannabis use by PWH was associated with reduced pro-inflammatory and oxidative stress gene expression and increased anti-inflammatory, β-oxidation, and brain-derived neurotropic factor gene expression (Figure 4)

Gene expression analysis of MDMs, using RNA sequencing and qPCR, revealed significant differences influenced by HIV status and cannabis use.

Compared to cannabis-naïve/low PWoH, daily cannabis-using PWoH showed minimal changes in gene expression. In contrast, MDMs from cannabis-naïve/low PWH exhibited a pro-inflammatory profile, with higher expression of genes such as IL-1α, IL-1β, IL-18, MARCO, and FPR2, and lower expression of anti-inflammatory genes, including IL-4, IL-10, CD163, GPR18, and CD200R1 (Figure 4, A – H). This profile was reversed in MDMs from PWH with moderate or daily cannabis use, which showed an anti-inflammatory phenotype (Figure 4, A–H).

Consistent with the observation of higher mitochondrial numbers in PWH who use cannabis, gene expression analysis showed lower levels of mRNA associated with oxidative stress. Specifically, RNA sequencing showed that genes related to oxidative stress, such as HIF1α, GPX3, and SOD1, were higher in cannabis-naïve/low PWH but downregulated in PWH with moderate and daily cannabis use (Figure 4, I–L).

Furthermore, a more active β-oxidation process was evident in MDMs from PWH who were moderate and daily users, with the upregulation of several key β-oxidation genes (CPT1A, ACADVL, ACAD10, ACAD11) (Figure 4, M–P).

Finally, BDNF expression followed a notable pattern. While it was downregulated in cannabis-naïve/low PWH, it was upregulated in both daily cannabis-using PWoH and PWH with moderate and daily cannabis use (Figure 4, Q–T).

Thus, these findings suggest that cannabis use by PWH is associated with increased anti-inflammatory cytokines, β-oxidation, and BDNF gene expression, and decreased oxidative stress gene expression in MDMs.

*Cannabis use by PWH was associated with lower plasma levels of growth differentiation factor 15, soluble triggering receptor expressed on myeloid cells 2, and increased mature BDNF/precursor BDNF ratios that correlated with better cognition (Figure 5) Inflammatory Markers (GDF15)*:

**Figure 5.**
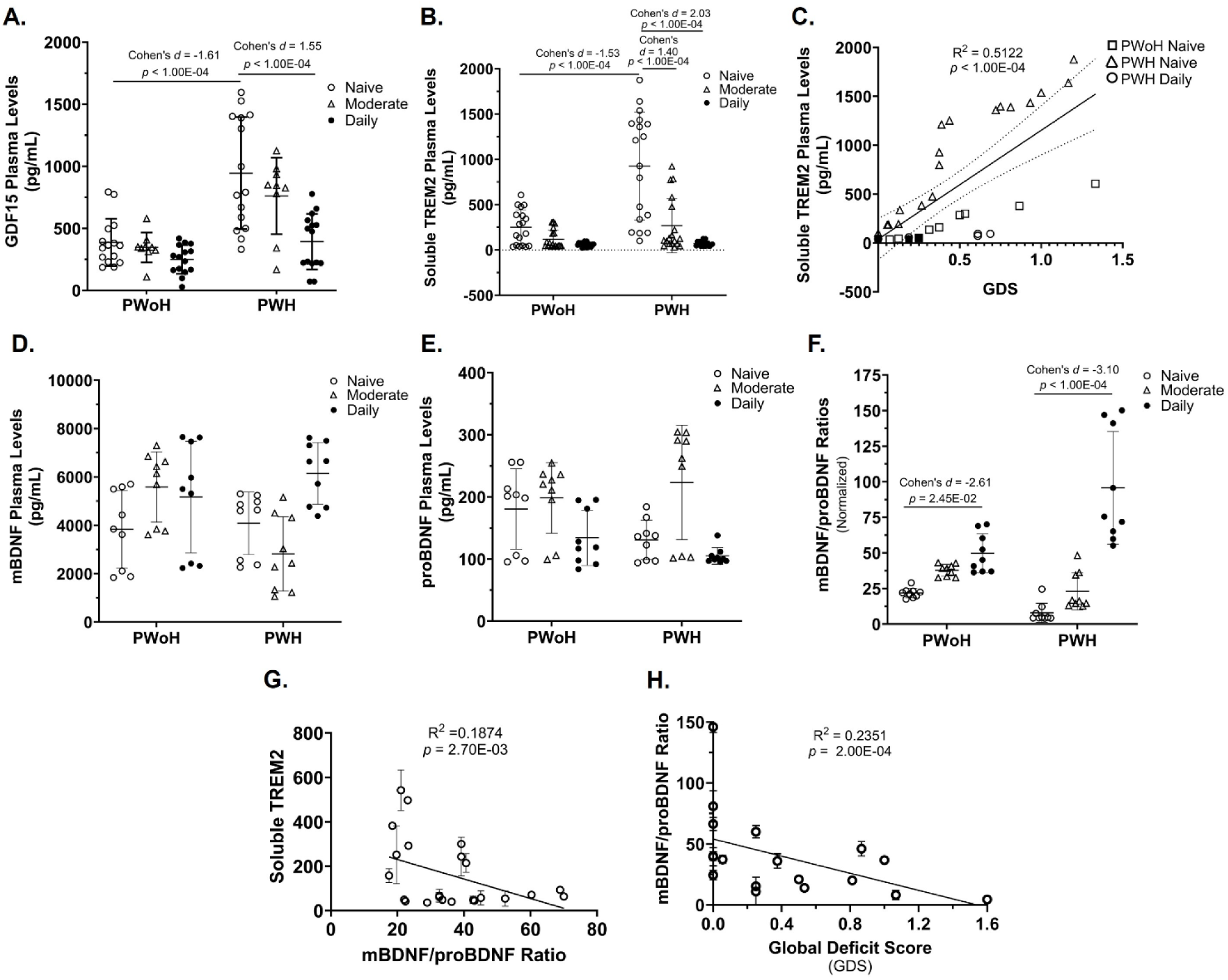
Cannabis use by PWH was associated with lower plasma levels of growth differentiation factor 15, soluble triggering receptor expressed on myeloid cells 2, and increased mature BDNF/precursor BDNF ratios that correlated with better cognition. (**A)** Quantification of GDF15 plasma levels via ELISA (Intra-assay CV = 5.6%, LLOQ: 1.0 pg/mL). (**B**) Quantification of sTREM2 plasma levels via ELISA (Intra-assay 1 CV = 4.6%, Intra-assay 2 CV = 5.7%, Inter-assay CV = 5.2%, LLOQ: 0.7 pg/mL). (**C**) Correlation of plasma sTREM2 levels with Global Deficit Score (GDS). (**D – F**) Quantification of mBDNF (Intra-assay CV = 5.3%, LLOQ: 200 pg/mL) and proBDNF (Intra-assay CV = 8.6%, LLOQ: 30 pg/mL) plasma levels via ELISA and calculation of the mBDNF/proBDNF ratios. (**G**) Correlation of mBDNF/proBDNF ratios with sTREM2 levels. (**H**) Correlation of mBDNF/proBDNF ratios with GDS. mBDNF: mature BDNF; proBDNF: precursor BDNF; GDF15: Growth Differentiation Factor 15; TREM2: Triggering Receptor Expressed on Myeloid Cells 2; LLOQ: Lower Limit of Quantitation. *Statistics*: (A-E, G-I) Two-way ANOVA with Tukey’s Multiple Comparisons Test. Simple Linear Regression Analysis. Confidence intervals and effect sizes: GDF15 PWoH Naïve/Low vs PWH Naïve/Low: 95% CI [-974.09, -379.3], *d* = -1.61 and effect-size *r* = -0.63; GDF15 PWH Naïve/Low vs PWH Daily: 95% CI [572.39, 1138.76], *d* = 1.55 and effect-size *r* = 0.61; sTREM2 PWoH Naïve/Low vs PWH Naïve/Low: 95% CI [-790.59, -325.05], *d* = -1.53 and effect-size *r* = -0.61; sTREM2 PWH Naïve/Low vs PWH Daily: 95% CI [311.79, 790.6], *d* = 2.03 and effect-size *r* = 0.71; mBDNF/proBDNF Ratios PWoH Naïve/Low vs PWoH Daily: 95% CI [10.39, 17.59], *d* = 2.61 and effect-size *r* = 0.80; mBDNF/proBDNF Ratios PWH Naïve/Low vs PWH Daily: 95% CI [-106.9, -68.77], *d* = -3.10 and effect-size *r* = - 0.84; (A) n = 3 donors with 3 biological replicates per group; (B) n = 6 donors with 3 biological replicates per group; (C) n = 6 donors with 3 biological replicates per group; (D – F) n = 3 donors with 3 biological replicates per group; (G) n = 3 donors with 3 biological replicates per group; (H) n = 3 donors with 3 biological replicates per group, the mBDNF/proBDNF ratio was plotted with each donor’s GDS score.

A significant interaction was observed between HIV status and frequency of cannabis use concerning GDF15 plasma levels (F(2, 72) = 4.604, P = 0.0131). This finding indicates that the effect of HIV status on GDF15 plasma levels varies with the frequency of cannabis use (Figure 5A).

Specifically, cannabis-naïve/low PWH had significantly higher plasma GDF15 levels compared to cannabis-naïve/low PWoH (*p* < 1.00 × 10^-4^, 95% CI [-974.09, -379.3]). In contrast, daily cannabis use by PWH was associated with a significant reduction in GDF15 relative to cannabis-naïve/low PWH (*p* < 1.00 × 10^-4^, 95% CI [572.39, 1138.76]) (Figure 5A).

### Neuroinflammatory and Neurotrophic Markers (sTREM2, BDNF)

A statistically significant interaction was observed between HIV status and frequency of cannabis use concerning plasma sTREM2 levels (*F*(2, 102) = 13.60, *P* < 0.0001). This indicates that the effect of HIV status on sTREM2 plasma levels varies depending on cannabis use frequency (Figure 5B).

Specifically, cannabis-naïve/low PWH had significantly higher plasma sTREM2 levels compared to cannabis-naïve/low PWoH (*p* < 1.00 x 10^-4^, 95% CI [-790.59, -325.05]). However, sTREM2 levels were significantly reduced in PWH who were moderate or daily cannabis users (*p* < 1.00 x 10^-4^, 95% CI [311.79, 790.6]) (Figure 5B).

Moreover, sTREM2 levels correlated positively with GDS (*p* < 1.00 x 10^-4^, 95% CI [740.9, 1487]), indicating that higher sTREM2 levels are associated with greater neurocognitive deficits (Figure 5C).

A statistically significant interaction was observed between HIV status and frequency of cannabis use concerning plasma mBDNF/proBDNF ratios (*F*(2, 48) = 16.41, *P* < 0.0001). The effect of HIV status on these ratios changed depending on cannabis use frequency (Figure 5F).

The mBDNF/proBDNF ratio was significantly higher in daily cannabis-using PWoH compared to cannabis-naïve/low PWoH (*p* = 2.45 × 10^-2^, 95% CI [10.39, 17.59]).

Similarly, the ratio was significantly higher in daily cannabis-using PWH compared to cannabis-naïve/low PWH (*p* < 1.00 × 10^-4^, 95% CI [-106.9, -68.77]) (Figure 5, D – F). Furthermore, sTREM2 plasma levels correlated significantly with mBDNF/proBDNF ratios (*p* = 2.70 × 10^-3^, 95% CI [-7.173, -1.614]) (Figure 5, G), and the mBDNF/proBDNF ratio correlated significantly with GDS (*p* = 2.00 × 10^-4^, 95% CI [-52.68, -17.47]) (Figure 5, H). Thus, these findings suggest that in PWH, cannabis use is associated with reduced inflammation and increased mBDNF/proBDNF ratios, which correlate with better cognitive function.

## Discussion

This study demonstrates that cannabis use, particularly daily use, by PWH is associated with a distinct immunometabolic phenotype in monocyte-derived macrophages (MDMs) characterized by evidence of low glycolysis, high oxidative phosphorylation (OXPHOS), and elevated anti-inflammatory gene expression. This cellular phenotype corresponds with better neurocognitive performance and lower plasma inflammatory biomarkers in PWH using cannabis (26,26–31). These findings are particularly significant given that MDMs are critical mediators of HIV-associated comorbidities: they are among the first cells infected during viral transmission, traffic HIV to the brain, and produce inflammatory cytokines that affect bystander cells and the systemic inflammatory milieu (55–57). Understanding MDM phenotype in PWH and its relationship to cognitive function is therefore critically important for identifying therapies against HIV-associated neurocognitive impairment (NCI). This is the first study to report associations between an anti-inflammatory immunometabolic profile in patient-derived MDMs to higher average cognitive outcomes in PWH using cannabis, building on prior work from our group showing cannabis use is associated with changes in TREM2 expression and NLRP3 inflammasome-related gene expression in MDMs (33,58).

A metabolic shift toward glycolysis typically drives inflammatory responses in macrophages and other myeloid lineage cells (59). While some studies of HIV-infected MDMs show shifts toward either OXPHOS or glycolysis (15,60–62), this study examined uninfected bystander MDMs from PWH on ART, as the majority of circulating PBMCs are not productively infected (63–65). The signal reflecting high OXPHOS and relatively low glycolysis observed in MDMs from PWH using cannabis is consistent with an anti-inflammatory, tissue-reparative phenotype (66). Such immunometabolic reprogramming is implicated in processes beyond energy production to influence biosynthetic processes, redox balance, reactive oxygen species levels, mitochondrial density, and cellular functions including proliferation, differentiation, and genomic integrity (67,68). As critical mediators of innate immunity, macrophages orchestrate both inflammatory and reparative responses to maintain tissue homeostasis (69–72). The pro-OXPHOS gene expression associated with cannabis use suggests MDMs can contribute to resolving chronic inflammation in PWH. This is further supported by low levels of mRNA associated with oxidative stress and high levels of mRNA for the reparative factor TGF-β2 (73,74), consistent with enhanced tissue repair and inflammation resolution (75–78). Critically, relatively high BDNF mRNA expression in MDMs from PWH using cannabis suggests enhanced production of secreted BDNF, which promotes neuronal growth, synaptic plasticity, and cognitive function (79–81), collectively consistent with a neuroprotective MDM phenotype that may mitigate NCI in PWH. However, longitudinal studies and clinical trials using larger cohorts with rigorous inclusion and exclusion criteria are necessary to better test the effect of cannabis use on MDM phenotype and brain function in PWH.

The effects of cannabis use on neurocognitive outcomes is likely contextual, depending upon disease state, population, and individual characteristics, among other factors. Prior studies in PWH report mixed cognitive findings, with some showing associations consistent with reduced inflammation and better cognitive outcomes and others reporting null or adverse effects, highlighting substantial variability across cohorts and study designs (82–87). However, studies of neurocognitive outcomes associated with neutral or adverse effects in people without HIV (PWoH) reflecting fundamental differences in underlying pathophysiology (88–93). One plausible explanation for these divergent observations is the distinct inflammatory and metabolic baseline associated with chronic HIV infection, which is characterized by persistent immune activation, mitochondrial injury, and oxidative stress even under effective antiretroviral therapy (58,94,95). In this pathological context, the anti-inflammatory properties of cannabinoids may partially mitigate HIV-associated neuroinflammatory processes and their neurotoxic effects (26,27), with benefits most pronounced in cognitive domains disproportionately affected by HIV such as verbal fluency, learning, and motor function (96). Conversely, in healthy individuals without chronic neuroinflammation, acute or chronic THC exposure can impair mitochondrial respiration via mitochondrial CB1 receptors and disrupt glutamatergic and GABAergic signaling critical for synaptic plasticity and memory (33,97–102). Within the HIV pathological environment, cannabis-associated anti-inflammatory effects, particularly suppression of NLRP3 inflammasome activity (95,103), may preserve mitochondrial integrity against ongoing inflammatory damage, and evidence in PWH suggests an association with normalization of glutamate concentrations and stabilization of GABAergic cortical dynamics (104–106). These effects represent relative improvements compared to non-using PWH rather than enhancement beyond healthy baselines. However, the current finding that THC induced higher levels of mitochondrial activity in MDMs generated from PWH using cannabis with better cognitive scores is consistent with a role of cannabis in promoting an anti-inflammatory phenotype in these individuals. Strikingly, this relationship was not apparent in THC-treated MDMs from PWH in the cannabis naïve group, which could indicate that cannabis use primes MDMs for THC-induced increases in mitochondrial activity. Conversely, this finding could reflect a difference in how some people respond to cannabis use, and this may contribute to personal choices to use cannabis recreationally or medically. These differences in relationship could also indicate that PWH using cannabis may be more cognitively resilient and not that cannabis is causing the neuroprotection. A clinical trial would be needed to test this hypothesis. However, the positive effects of cannabis use in some PWH are consistent with studies showing that cannabinoids can be neuroprotective and anti-inflammatory in rodent models for neurodegenerative diseases and even in a clinical trial of people exhibiting Alzheimer’s like symptoms (107–109). This framework positions cannabis as a potential pathology-specific modulator of HIV-driven neuroinflammation rather than a generalized cognitive enhancer and observed effects should be interpreted cautiously as domain-specific and context-dependent rather than implying global cognitive benefit (31).

Specific doses of cannabis or combinations of phytocannabinoids may be more beneficial for some people than others, which is consistent with a previously described hormetic effect of cannabis (110–112). Cannabis exposure itself is heterogeneous, encompassing variability in frequency, dose, potency, cannabinoid composition (including THC:CBD ratios), and route of administration, and these factors may differentially modulate immune and metabolic pathways depending on the presence or absence of HIV-related inflammation. In the present study, the daily-use category includes individuals reporting once-daily use as well as those using multiple times per day. Future research in larger daily use samples should examine whether lighter versus heavier daily use may differentially alter immunometabolic or neurocognitive profiles. Moreover, neurocognitive outcomes in PWH are influenced by numerous factors (e.g., comorbid medical and psychiatric conditions, vascular risk, polysubstance use, and lifestyle variables) that were not comprehensively modeled in this cross-sectional study. Individual differences withing populations of people with neurodegenerative diseases may deem some people more amenable to cannabis or cannabinoid-based therapies that others (111). Accordingly, these findings should be interpreted as associative and context-dependent. While cannabis-related immunometabolic modulation may contribute to differences in cognitive vulnerability in PWH, future studies with larger samples and more granular and/or objective measures of cannabis exposure may improve our understanding of these relationships.

The endocannabinoids system is increasingly being investigated as a therapeutic target against diseases with an inflammatory component (113). The observed effects in the current study may reflect engagement of the endocannabinoid system (ECS), particularly CB2 receptors, which are primarily expressed on MDMs alongside minimal CB1 expression (114,115). CB2 receptor activation mediates anti-inflammatory activity, influences cellular metabolism, and promotes mitochondrial function (116–118). THC, the predominant cannabinoid in cannabis, is a partial agonist for CB1 and CB2 (119).

Selective CB2 agonists such as JWH015 induce shifts from pro-inflammatory to homeostatic phenotypes and demonstrate neuroprotection in rodent neurodegeneration models (120–124). Prior work demonstrates that cannabis use is associated with relatively high levels of TREM2-related transcripts in MDMs (33), and cannabinoids suppresses pro-inflammatory IL-1β secretion, and can reverse HIV-induced metabolic reprogramming that disrupts synaptic plasticity (125). Cannabis use is also associated with reductions in activated and inflammatory immune cell frequencies and alterations in viral reservoir dynamics in PWH on ART (126,127). These findings are consistent with the hypothesis that cannabis promotes an anti-inflammatory immunometabolic phenotype in PWH MDMs primarily through CB2-mediated pathways (27,128,129), highlighting the potential therapeutic utility of non-psychoactive CB2-selective agonists for HIV-associated NCI.

Plasma biomarkers provide robust systemic validation of the cellular findings (130). Cannabis-using PWH exhibited lower levels of GDF15, a biomarker of oxidative stress, inflammation, and mitochondrial dysfunction that is elevated in neurodegenerative disease and associated with HIV-associated NCI (44,131), suggesting cannabis use may be associated with reduced systemic inflammation and improved mitochondrial health. Reduced soluble TREM2 (sTREM2) in cannabis users is particularly significant given its positive correlation with global neurocognitive deficits and elevated CSF levels in HIV-associated neurocognitive impairment (33,132–134); mechanistically, TREM2 regulates metabolic homeostasis in microglia, promoting mitophagy and reducing neuroinflammation (122–124). Although sTREM2 is not yet an established plasma biomarker for NCI in PWH, these findings underscore its potential in this context. Daily cannabis users also demonstrated a higher plasma mBDNF/proBDNF ratio, a key neuroplasticity indicator inversely correlated with global neurocognitive deficits (130,132). Collectively, these converging biomarkers—reduced inflammatory and stress markers (GDF15, sTREM2) alongside increased neuroplasticity indicators (mBDNF/proBDNF)—provide strong systemic evidence complementing the cellular and molecular data, suggesting cannabis use may mitigate inflammation and support neurocognitive function in PWH. Importantly, better controlled longitudinal studies and clinical trials will be needed to solidify these associations.

Several important limitations of this study must be acknowledged. The cross-sectional design and limited sample size preclude causal inference, and reliance on self-reported cannabis use requires validation with objective plasma cannabinoid measurements. The respirometry data showed high inter-individual variability and a small sample size, rendering these findings prone to effect size inflation and Type II errors; they should be viewed as exploratory proof-of-concept data requiring validation in larger, more diverse cohorts. The sample size precludes controlling for important confounding variables, including polysubstance use (135–137), psychiatric comorbidities, variable ART regimens with differing CNS penetration and neurotoxicity profiles (133,134,138,139), and route of cannabis administration. Based on the current findings, we cannot rule out the possibility that cannabis-using PWH are inherently more cognitively resilient through mechanisms not investigated in this study. The current studies did not assess the phenotype of other important immune cells that may be mediating the effects of cannabis in PWH and this should be considered in future studies. Future research requires longitudinal designs with precise characterization of cannabis use including dose, potency, and THC:CBD ratios, along with control for key confounders and ultimately randomized controlled trials to rigorously test whether cannabis causally promotes neuroprotection through immunometabolic reprogramming.

Despite these limitations, this work provides evidence of a mechanistic link through which cannabis may mitigate HIV-associated NCI by immunometabolically reprogramming MDMs toward an anti-inflammatory, neuroprotective phenotype (27, 31). These findings underscore the therapeutic potential of targeting immunometabolic pathways to reduce inflammation and support cognition in PWH and cautiously raise the possibility of adjunctive cannabinoid-based therapies—including non-psychoactive CB2-selective agonists—for HIV-associated NCI. The inherent risks of cannabis use, and the conflicting literature nonetheless necessitate careful interpretation and further large-scale, longitudinal investigation before any clinical recommendations can be made.

## Declarations

### Ethical Approval and Consent to Participate

Written informed consent to participate was received from all participants prior to inclusion in the study, and all human data associated with this study were performed in accordance with the Declaration of Helsinki with ethics approval by the UC San Diego Institutional Review Board Administration (IRB) Reference # 808184.

### Consent for Publication

Written informed consent for publication was received from all participants prior to inclusion in the study.

## Availability of Data and Materials

The datasets supporting the conclusions of this article are included within the article, and the RNAseq data was published in the GEO database and is accessible under accession number GSE285294 and available at <u>GEO Accession viewer</u>. The gene expression analysis code was submitted to the international open-access repository Dryad.

## Author Contributions

MKF, PWH, SL, RJE, JI, and JAF all participated in experimental design. PWH and JAF wrote the initial manuscript text. JAF, JI, JS, MCGM, RJE, JS, PWH, and MKF edited the original manuscript. PWH and JAF wrote the final manuscript version. MKF, AEL, GF, and AAL collected the blood samples. MKF, PWH, AEL, BLA, AB, MS, JIM, KCW, DDH, GF, EGS, and AAL conducted experiments, analyzed data, and made figures. MKF, PWH, RJI, ST, MCGM, JS, JI, and JAF analyzed data, formatted figures and improved data analyses.

## Acknowledgements

We acknowledge the UC San Diego IGM Genomics Center because this publication includes data generated from the IGM by utilizing an Illumina NovaSeq X Plus that was purchased with funding from the National Institutes of Health SIG grant (#S10 OD026929); We acknowledge the HIV Neurobehavioral Research Program (HNRP), UCSD (La Jolla, CA) for their recruitment of donors as well as their efforts that made this study possible.

## Legends

### Graphical Abstract

Cannabis use by people with HIV (PWH) is associated with neuroprotective and anti-inflammatory effects, mediated in part by reprogramming the immunometabolism of monocyte-derived macrophages (MDMs). This shift involves a transition from a pro-inflammatory, glycolytic phenotype toward a more anti-inflammatory state characterized by oxidative phosphorylation and altered cytokine expression.

In cannabis-Naïve/Low PWH, MDMs exhibit a pro-inflammatory profile that promotes neuroinflammation and mitochondrial damage, resulting in a reduced number of mitochondria. Chronic neuroinflammation contributes to neurodegeneration and the neurocognitive impairments (NCI) that define HIV-Associated Neurocognitive Disorders (HAND). This inflammatory and neurodegeneration profile is reflected by plasma immunometabolic biomarkers including reduced mature BDNF (mBDNF), and elevated GDF15 and soluble TREM2 (sTREM2) levels.

Conversely, cannabis use in PWH induces an anti-inflammatory MDM profile that promotes reduced neuroinflammation with increased mitochondrial biogenesis. The reduced neuroinflammation leads to improved neuronal function and reduced NCI, resulting in improved memory, motor and verbal skills, and executive learning. This neuroprotective effect is corroborated by corresponding plasma biomarkers showing increased mBDNF levels, and decreased GDF15 and sTREM2 levels. Collectively, these findings suggest that cannabis use by PWH appears to mitigate neuroinflammation and NCI, with the immunometabolic reprogramming of MDMs likely playing a key mechanistic role.

### Graphical Methods

The HIV Neurobehavioral Research Program (HNRP) donor cohort consisted of people without HIV (PWoH) and people with HIV (PWH) who were either cannabis naïve/low, moderate cannabis users, or daily cannabis users. Peripheral blood mononuclear cells (PBMCs) were isolated from donor blood samples, plated, and allowed to differentiate for two weeks into monocyte-derived macrophages (MDMs) before being tested or treated with THC and then tested. At the same time plasma samples were isolated from each donor and stored for testing. At that point specific dyes were added, and different parameters were measured such as on the CX5 CellInsight Imager or an ELISA and the data was then analyzed.

